# A nonsense mutant of the hepatitis B virus large S protein antagonizes multiple tumor suppressor pathways through c-Jun activation domain-binding protein1

**DOI:** 10.1101/477919

**Authors:** Shu-Yi Chiu, Hsiang-Ju Chung, Ya-Ting Chen, Min-Syuan Huang, Chien-Chih Huang, Shiu-Feng Huang, Isao Matsuura

## Abstract

Hepatitis B virus (HBV) is a major cause of hepatocellular carcinoma (HCC). Previous studies have identified recurrent nonsense mutations in the HBV large S (LHBs) gene from the liver from HBV core antigen-positive HCC patients. These nonsense mutants have been shown to be oncogenic in mouse xenograft models using a mouse embryonic fibroblast cell line. Here, we expressed in a liver cell line Huh-7 a carboxy terminally truncated protein from a nonsense mutant of the LHBs gene, sW182* (stop codon at tryptophane-182). Although the sW182* protein appeared not to be very stable in the cultured liver cells, we confirmed that the protein can be highly expressed and retained for a prolonged period of time in the hepatocytes in the mouse liver, indicating its stable nature in the physiological condition.

In the Huh-7 cells, the sW182* mutant downregulated tumor suppressors p53 and Smad4. This downregulation was reversed by a proteasome inhibitor MG132, implying the involvement of proteasome-based protein degradation in the observed regulation of the tumor suppressors. On the other hand, we found that c-Jun activation domain-binding protein 1 (Jab1) physically interacts with the sW182*, but not wild-type LHBs. RNA interference (RNAi) of Jab1 restored the levels of the downregulated p53 and Smad4. The sW182* mutant inhibited the promoter activity of downstream target genes of the tumor suppressors. Consistently, Jab1 RNAi reversed the inhibition.

These results suggest that the LHBs nonsense mutant antagonizes the tumor suppressor pathways through Jab1 in the liver contributing to HCC development.

## Introduction

Hepatocellular carcinoma (HCC) is the most common primary malignancy of the liver and among the top causes of cancer death worldwide [1]. The high incidence of the disease is in Africa and Asia is attributed to endemic prevalence of hepatitis B (HBV) and C (HCV) viruses in the regions [1]. Chronic liver diseases such as cirrhosis caused by the viral infection often advance to HCC [2-5]. Thus it is generally accepted that HCC development by HBV infection is through the indirect effect of the virus that involves chronic liver injury [6]. In some cases, however, HCC can develop without cirrhosis, suggesting the viral factor(s) alone could be oncogenic [7]. Among the seven proteins encoded by four HBV genes, X protein encoded by X gene has been relatively well studied for its putative oncogenicity [8-12]. In addition to X protein, the large form of S protein (LHBs) encoded by Pre-S/S gene has been reported as a possible oncogenic factor. Mutant LHBs proteins with deletion in the Pre-S1 or Pre-S2 region have been identified in ground glass hepatocytes in late phase of chronic HBV infection [13,14]. The Pre-S2 mutant has been shown to regulate cell cycle-related proteins such as cyclin A and p27^Kip1^ [15,16], which may contribute to its putative oncogenic potential [17]. Recently, multiple nonsense mutations in S gene have been identified from cancerous parts of the HBV core antigen-positive (HBcAg(+)) HCC livers [18,19]. The detection of HBcAg implies the presence of viable, freely replicative HBV in the HCC liver tissues. Thus it is possible that the nonsense mutations may contribute to early stages of hepatocarcinogenesis [19]. The functional studies have shown that the nonsense mutations were indeed oncogenic [18-20]. In these functional studies, however, a mouse fibroblast cell line NIH3T3 was used as a model cell line. Since the cell line is unrelated to the liver, there are potential arguments about the relevance of the results. Therefore, in the current study we used Huh-7, a cell line of liver origin, to express LHBs sW182*, one of the three nonsense mutants with the highest oncogenic potential [19]. Probably because of the unstable nature of the sW182* protein in the liver cells, attempts to generate cell clones that express the mutant protein were unsuccessful. Nonetheless, using the transient expression system, we found that the LHBs sW182* protein can antagonize multiple tumor suppressor pathways involving p53 and Smad4. The mutant viral protein can directly interact with c-Jun activation domain-interacting protein 1 (Jab1), and Jab1 is necessary for downregulation of the tumor suppressors. Since oncogenic function of Jab1 has been recognized [21-23], it is likely that the LHBs nonsense mutant can contribute to hepatocarcinogenesis through Jab1.

## Materials and Methods

### Cell culture and transfection

HEK293T cells and Huh-7 cells were obtained from the repository of our Institute and maintained in DMEM, 10 % FBS in 5 % CO_2_ at 37°C. The LHBs and its nonsense mutant cDNA were cloned in pCDNA6-myc-His for C-terminal Myc-6xHis tag and pCMV2B for N-terminal Flag tag, respectively. For GST-pulldown assay, HEK293T cells were transfected with 9 μg pCMV2B-Flag-LHBs or the mutant in 100 mm plate using Lipofectamine (Thermo Fisher) following the manufacture’s instruction.

For the p53 analysis, Huh-7 cells (in 12-well plates) were co-transfected with 0.2 μg pCMV5-Flag-p53 and 1 μg pCDNA6-LHBs-myc-His or pCDNA6-LHBs (sW182*)-myc-His using Lipofectamine LTX (Thermo Fisher) following the manufacture’s instruction. For the Smad4 analysis, the cells were co-transfected with 0.6 μg pCDNA5-DPC4-HA, 25 ng pCMV5-TβRI (T204D)-His and 0.6 μg pCDNA6-LHBs-myc-His or pCDNA6-LHBs (sW182*)-myc-His. The pCDNA6 empty vector was used for a control for the viral protein expression.

Stealth RNAi duplexes against Jab1 were purchased from Invitrogen. The sequences of two oligos (forward) are (1) 5’-GGUGAUUGAUCCAACAAGAACAAU-3’ and (2) 5’-GAGUACCAGACUAUUCCACUUAAUA-3’. For Jab1 knock-down, Huh-7 cells were reverse transfected on seeding with the Jab1 RNAi duplexes by Lipofectamine 2000 (Thermo Fisher) following the manufacture’s instruction. After overnight the cells on the plate were further transfected with LHBs sW182* and p53 or Smad4/TβRI (T204D) constructs as above.

For the luciferase assay, the cells in 12-well plates were co-transfected with 0.4 μg reporter construct (human p21 (2.1kb)-Luc [24], 3TP-Lux [25], human Smad7-Luc [26]) and 0.5 μg LHBs constructs.

### Hydrodynamic injection

The LHBs gene fragment was used to replace the *BglII* to *NotI* region (containing CMV promoter) of pRc/CMV. Thus, the construct expressed the LHBs gene driven by its PreS1 viral promoter, not the CMV promoter. A stop codon was introduced in the wild-type gene by the site-directed mutagenesis to generate the sW182* construct.

Ten micrograms of either plasmid DNA was injected into the tail veins of C57BL/6 mice (male, 6 to 7 weeks old) in a volume of PBS equivalent to 8 % of the mouse body weight. The total volume was delivered within 5 sec. The blood was taken periodically starting 1 day post injection for the serum level of the viral protein. The mice were sacrificed after 24 days post injection and the liver tissue was analyzed by IHC for the HBsAg using the LHBs antibody (Viostat).

All animals in these experiments received human care. The protocol complied with our institution’s guideline and was approved by Animal Research Ethics Committee at National Health Research Institutes, Taiwan (IACUC approval No.: NHRI-IACUC-101124).

### GST-pulldown, immunoprecipitation and Western blot

Jab1 cDNA was cloned in a plasmid pGEX-KG and expressed as a GST-fusion protein. GST-Jab1- and control GST-beads were prepared as described previously [27]. Cell lysate was prepared from the transfected HEK293T cells (Flag-LHBs or sW182* mutant) in protein lysis buffer (0.1 % Na-deoxtcholate, 10 mM Tris, pH7.8, 1 mM EDTA, 0.2 mM EGTA, 1 % Np-40, 140 mM NaCl) supplemented with 1mM DTT, 25 ng/μl RNaseA, 1x cOmplete protease inhibitor cocktail (Merk). The call lysate (400 μg) and 15 μl beads were mixed and incubated at 4oC for 2 h. The beads were washed 5 x with lysis buffer containing 1 mM DTT, 25 ng/μl RNaseA, 1 mM benzamidine, 1 mM PMSF. The proteins on the beads were eluted with SDS sample buffer and a half of the sample was loaded on SDS PAGE followed by Western blot with Flag antibody (M2) (Sigma).

For co-immunoprecipitation of LHBs and Jab1 protein, 750 μg lysate from HEK293 cells transfected with pCMV2B-Flag-Jab1 and pCDNA6-LHBs-Myc-6xHis or pCDNA6-LHBs (sW182*)-Myc-6xHis was mixed with Myc-agarose beads (Sigma) for 4 h. After washing, the beads were eluted with SDS sample buffer. The eluted Jab1 protein was detected by Flag antibody.

Huh-7 cell lysate was also prepared as above. When MG132 was used, the transfected cells were treated with the drug (10 μM) for 6 h before being harvested. The antibodies used for the Western blot are: 6xHis, Genetex; Hemagglutinin (HA) (16B12), COVANCE; Myc (9E10), GeneTex; β-actin, Santa Cruz; Jab1 2A10, Gene Tex.

### Luciferase reporter assay

Cells were harvested 48 h post transfection and subjected to the luciferase assay following Muthumani’s protocol with the protein concentration of the lysate as a reference [28]. Each assay was performed at least three times with triplicates.

### Statistical analysis

Experiments were repeated at least three times, and representative results were shown. All data are presented as means ± S.E. T-test was used to calculate the significance of the data. A value of *P* < 0.05 was considered statistically significant and *P*-value <0.05 was marked with (*), < 0.01 with (**) or <0.001 with (***).

## Results

### Expression of the LHBs sW182* truncation mutant in Huh-7 cells

Lee et al. and we have recently reported that a nonsense mutant of LHBs, sW182* (Fig. 1A), has an oncogenic potential, as nude mice xenografted with mouse embryonic fibroblasts (NIH3T3) transfected by the mutant showed strong tumorigenicity [18-20].

**Fig. 1.**
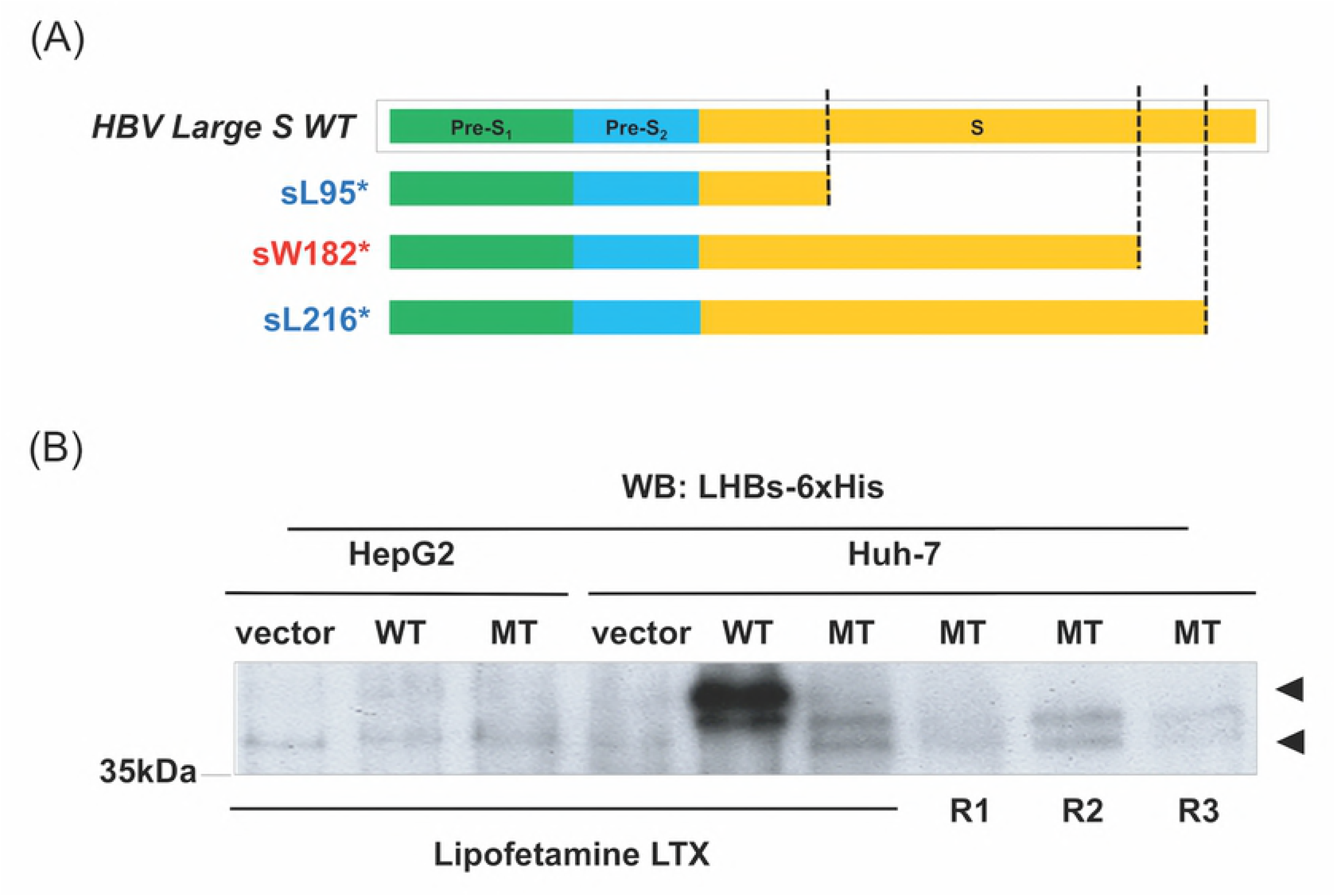
Expression of the LHBs sW182* truncation mutant in cultured liver cells. (A) Schematic diagram of the LHBs protein and its truncation mutants identified from the liver of HBV-related HCC [19]. The truncation mutants were named after the stop codon at the corresponding site for each amino acid. A nonsense mutation at the codons for L95, W182 and L216 resulted in sL95*, sW182* and sL216*, respectively. (B) HepG2 and Huh-7 cells were transfected with control vector, 6xHis-tagged wild-type (full length) LHBs (WT) or sW182* mutant (MT) using different transfection reagents Lipofectamine LTX, R1, 2, 3). The protein was detected by Western blot with the 6xHis antibody. The arrowheads indicate the position of the wild-type and mutant protein on the blot.

Since the cell line used in these studies was unrelated to the liver, however, it is necessary and relevant to attest its potential oncogenic mechanisms in liver cells. For this purpose we attempted to stably express the sW182* protein in liver cell lines. Inauspiciously, we failed to stably express the mutant protein in any of the liver cell lines we tested (HepG2, Huh-7, Tong and HA22T). We noted later that the sW182* protein cannot be expressed very well, even transiently, in the cultured liver cells. This is probably because of its instability in the liver cells. Fig. 1B shows expression of the wild-type LHBs and the sW182* truncation mutant in the transiently transfected liver cell lines (HepG2 and Huh-7) using different transfection reagents. In this condition, we failed to express the proteins in HepG2 cells, but both wild-type and the truncation mutant sW182* were expressed in Huh-7 cells. We noted that the expression level of the mutant was significantly lower than that of the wild-type protein, consistent with the notion of unstable trait of the mutant protein.

### Prolonged expression and retention of the LHBs sW182* truncation mutant in the mouse hepatocytes in vivo

The unstable nature of the LHBs sW182* mutant in the cultured liver cells raised an argument about its potential role in liver carcinogenesis. To test whether the truncation mutant can be expressed in the liver in vivo, we hydrodynamically injected mice with the expression construct for the wild-type LHBs or sW182* mutant under a hepatocyte-specific PreS1 promoter. After injection, blood was collected for the LHBs protein in the serum by HBsAg assay. The wild-type LHBs was detected at 1 day post injection (Table 1).

**Table 1.** LHBs levels in blood of hydrodynamically injected mice

| HBsAg Assay<sup>a</sup> |  |  |  |  |  |  |
| --- | --- | --- | --- | --- | --- | --- |
|  | Serum ID | Day 1<br>(IU/ mL) | Day 3<br>(IU/ mL) | Day 5<br>(IU/ mL) | Day 10<br>(IU/ mL) | Day 24<br>(IU/ mL) |
| WT | m1 | 1300 | 11800 | 10420 | >1300 | nd |
|  | m2 | 1300 | 14930 | 13080 | >1300 | 550 |
|  | m3 | 1300 | 10620 | 9670 | >1300 | 1289 |
| sW182* | m4 | nd<sup>a</sup> | nd<sup>a</sup> | nd<sup>a</sup> | nd<sup>a</sup> | nd<sup>a</sup> |
|  | m5 |  |  |  |  |  |
|  | m6 |  |  |  |  |  |

Blood samples were taken on the indicated days post-injection and the LHBs proteins in plasma were quantitated by ELISA.
<sup>a</sup>Not detected (<5)

The level of the protein peaked at 3 days post injection and gradually declined through the last day of the experiment (24 days post injection). On the other hand, the sW182* mutant was not detected in the blood throughout the experiment. In contrast, the immunohistochemistry (IHC) analysis showed a relatively high expression of the sW182* mutant in the liver cells, whereas the wild-type protein was hardly detected (Fig. 2).

**Fig. 2.**
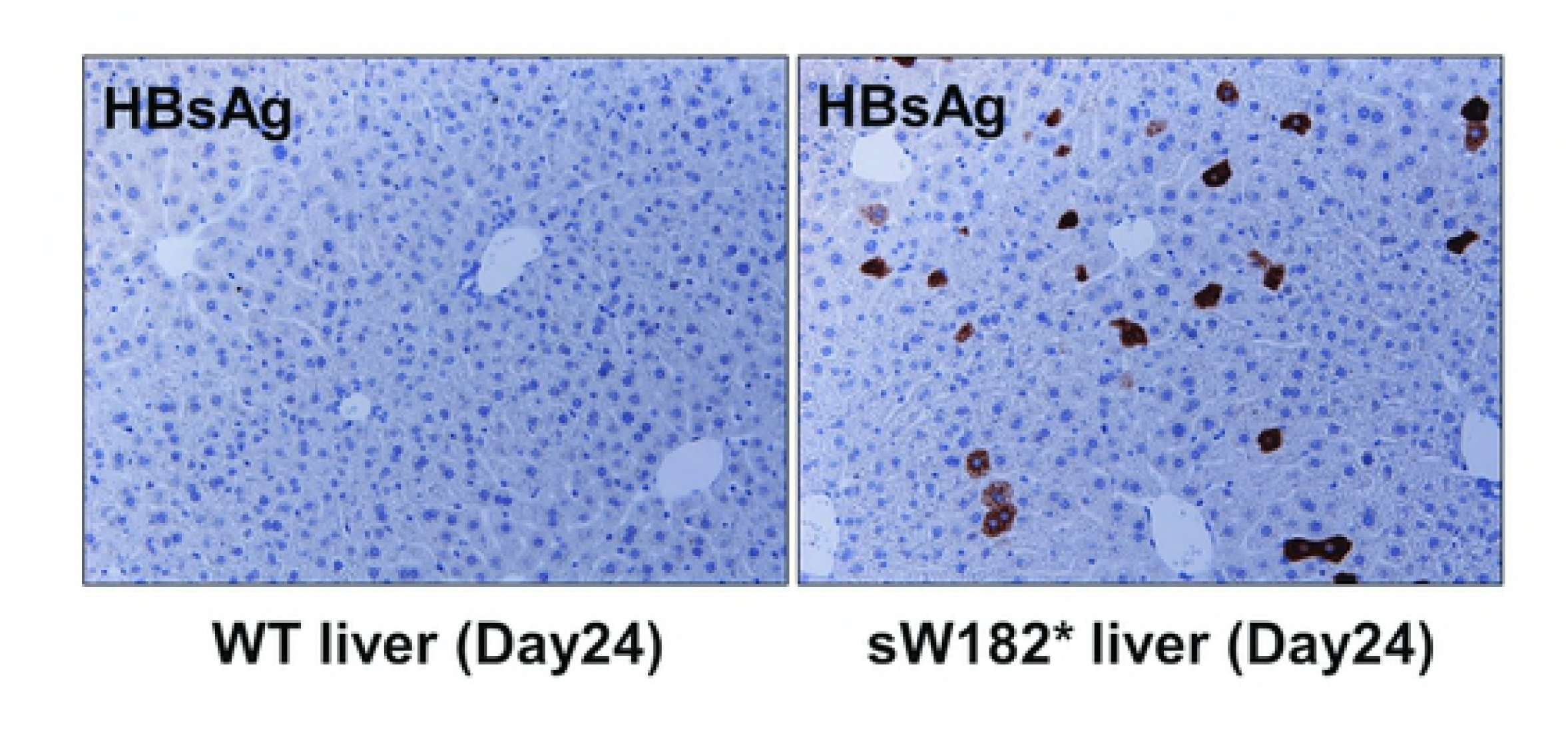
Expression of the LHBs sW182* truncation mutant in hepatocytes in the living mouse liver. Expression of LHBs proteins in hydrodynamically transfected hepatocytes in mouse liver. At 24 days post injection, mice were sacrificed and the liver was subjected to the IHC analysis with the LHBs antibody. Left panel, wild-type LHBs; right panel, sW182*.

This result demonstrated that the LHBs sW182* truncation mutant can be expressed and retained in the hepatocytes in the living liver for a prolonged period of time, supporting the relevance of the further functional investigation of the mutant using the cultured liver cells, regardless of its apparent instability in the system. The result also revealed a significant difference between the wild-type and truncation mutant in the liver retention in the physiological condition.

### The sW182* mutant, but not wild-type LHBs physically interacts with Jab1

c-Jun activation domain-binding protein 1 (Jab1) can promote cell proliferation and survival by influencing multiple pathways. Its activity can be enhanced by direct interaction with interacting proteins such as psoriasin (S100A7) [29]. Interestingly, similar type of interaction has been documented for a Pre-S2 mutant of LHBs, in which a part of Pre-S2 region of the protein is deleted [16]. Prompted by this study, we tested whether the LHBs sW182* truncation mutant could physically interact with Jab1. In the GST-pulldown assay GST-Jab1 precipitated the LHBs sW182*, but not wild-type protein, from the HEK293T cell lysate (We noted that the viral proteins can be well expressed in this cell line) (Fig. 3).

**Fig. 3.**
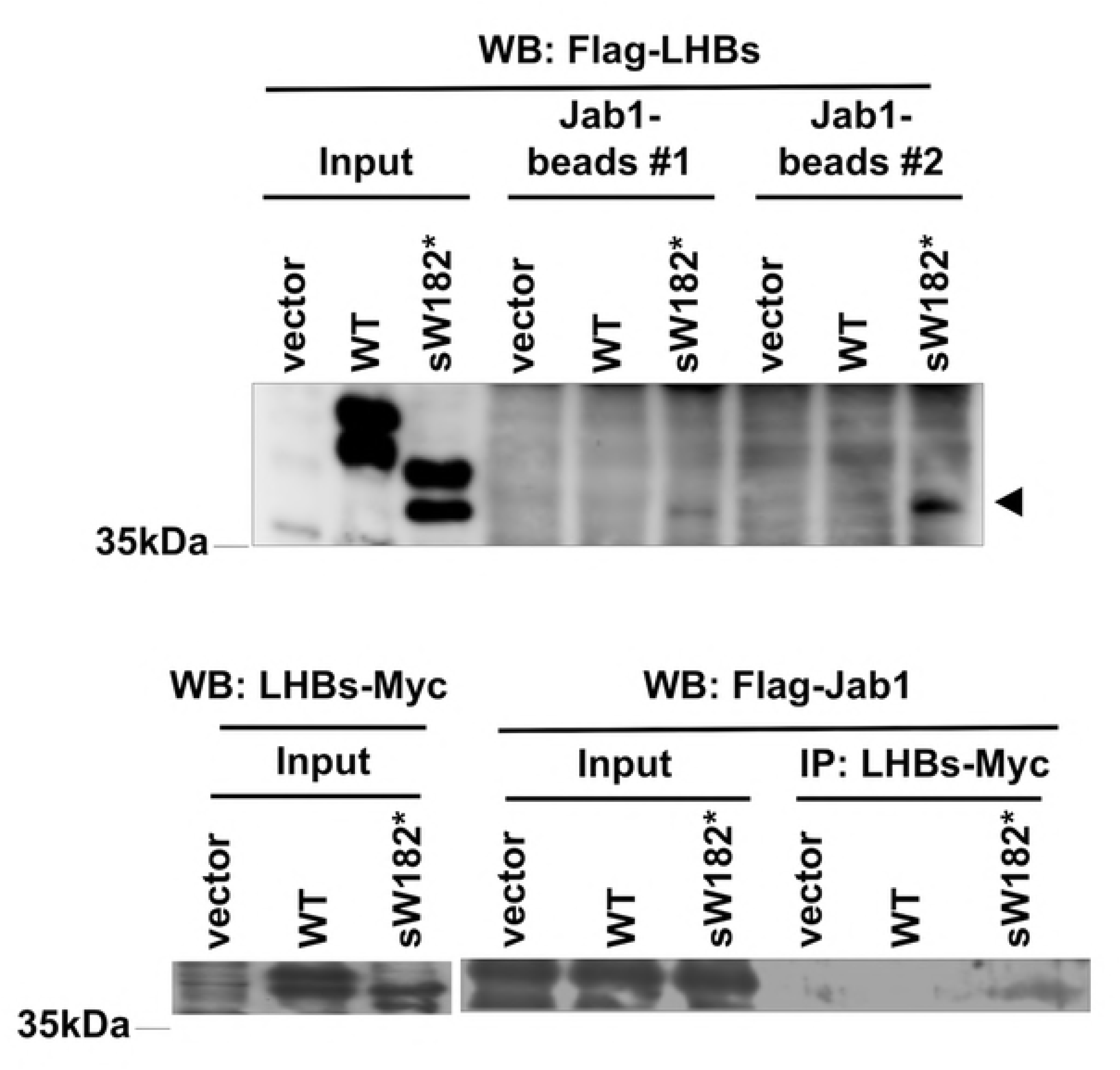
Physical interaction between Jab1 and the sW182* truncation mutant, but not wild-type LHBs. Upper panel: Flag-tagged wild-type LHBs or sW182* mutant expressed in HK293T cells was pulled down with GST-Jab1-beads. The precipitated proteins were subjected to Western blot using Flag antibody. Vector control was also included. Results with two independent preparations of GST-Jab1-deads are shown (#1 and #2). Lysate (6 %) used for the assay is shown as Input. The precipitated sW182* protein is indicated by an arrowhead.

Lower panel: Flag-tagged Jab1 and 6xHis-Myc-tagged wild-type LHBs or sW182* mutant were co-expressed in HK293T cells. Myc-tagged protein was precipitated by Myc antibody-conjugated agarose beads and Jab1 was detected by Flag antibody. Lysate (1.5 %) used for the assay is shown as Input.

Reciprocally, Jab1 was co-immunoprecipitated with the LHBs w182*, but not with wild-type protein, from the cell lysate (Fig. 3). The physical interaction strongly suggested a possible influence of the sW182* mutant on Jab1 function.

### The LHBs sW182* truncation mutant downregulates multiple tumor suppressor proteins

Jab1 has been reported to promote degradation of a number of proteins including multiple tumor suppressors. Based on the result of Fig. 3, we examined the effect of the sW182* truncation mutant on the levels of Jab1 target tumor suppressors, p53 and Smad4 [30-33]. We found that the sW182* mutant downregulates these two tumor suppressors (Fig. 4A,B).

**Fig. 4.**
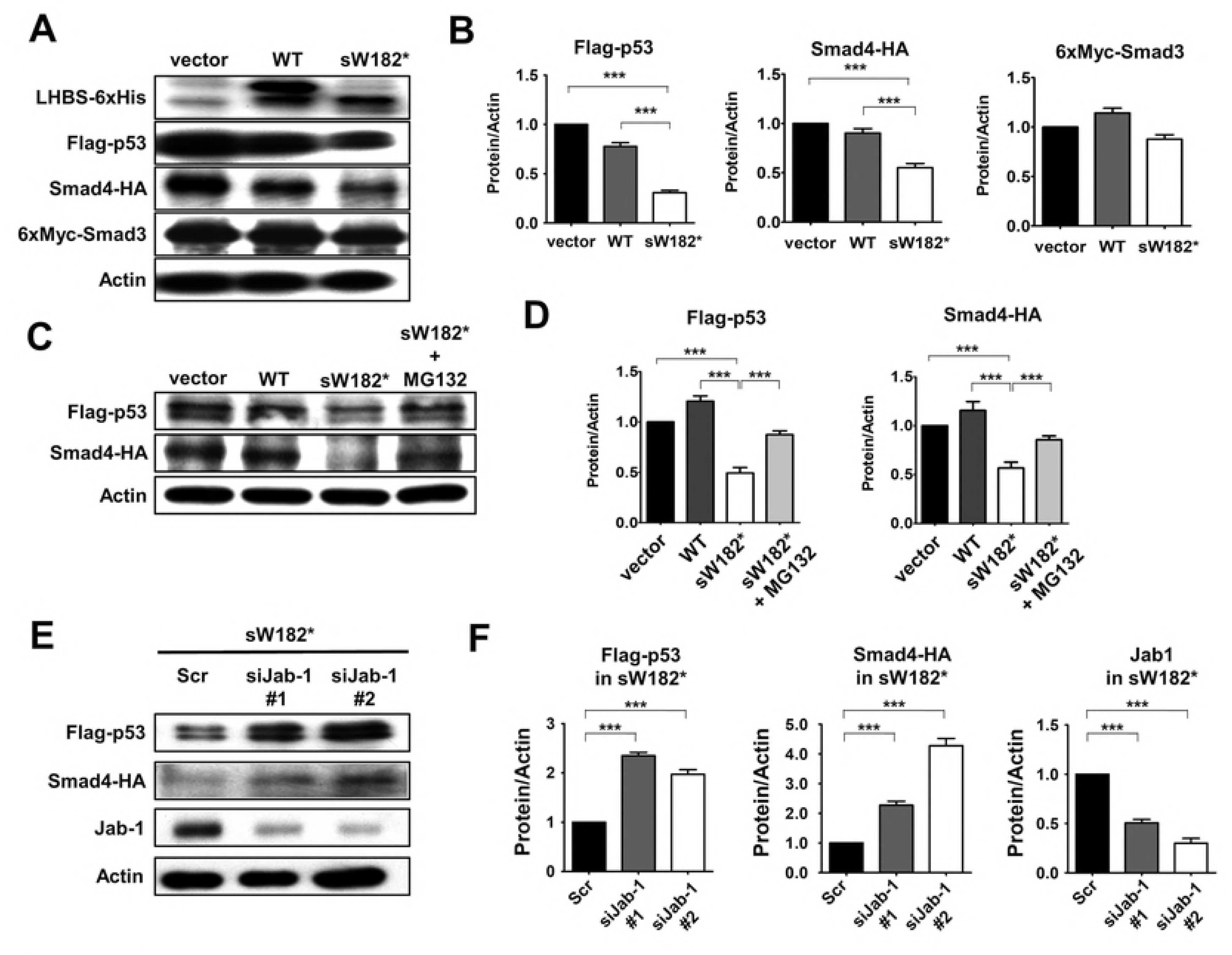
Proteasome- and Jab1-dependent downregulation of tumor suppressors p53 and Smad4 by the LHBs sW182* mutant. (A) Huh-7 cells were cotransfected with vector, wild-type LHBs or sW182* mutant, and Flag-p53, Smad4-HA or 6xMyc-Smad3. After 48 h the cells were harvested and the proteins were detected by Western blot using the corresponding antibody. Expression of the LHBs proteins and β-actin loading control are also shown. (B) The levels of p53, Smad4 and Smad3 in (A) and two other Western blots were quantitated using the β-actin loading control. (C) Huh-7 cells were transfected as in (A). Before harvesting, the cells were treated with MG132 for 6 h. (D) The levels of p53 and Smad4 in (C) and two other Western blots were quantitated using the β-actin loading control. (E) Huh-7 cells were transfected with siRNA (a control (Scr) and two for Jab1 (#1 and 2)), followed by the expression constructs for sW182* mutant and Flag-p53 or Smad4-HA. The levels of p53 and Smad4 were analyzed by Western blot. Depletion of Jab1 protein from the cells was also monitored. (F) The levels of p53 and Smad4 in (E) and two other Western blots were quantitated using the β-actin loading control. Results shown are presented as means ± S.E.

In contrast, the sW182* mutant had no effect on a non-Jab1 target Smad3 (Fig. 4A,B). Treatment of the sW182*-transfected cells with a proteasome inhibitor MG132 restored the levels of p53 and Smad4, indicating that the downregulation of these proteins are through the proteasome-based protein degradation (Fig. 4C,D). This observation was consistent with the possible involvement of Jab1 in protein degradation process, as Jab1 utilizes the proteasome for degradation of target proteins [30,33]. To test the involvement of Jab1 in the downregulation of the tumor suppressors by the LHBs sW182* mutant, we depleted the Jab1 from the liver cells by siRNA (Fig. 4E,F). As anticipated, the levels of p53 and Smad4 that had been downregulated by the sW182* mutant were restored in the Jab1-depleted cells (Fig. 4E,F). These results clearly demonstrated that the truncation mutant of the LHBs employs the Jab1-proteasome pathway to downregulate p53 and Smad4.

### The LHBs sW182* truncation mutant inhibits downstream genes of p53 and Smad4

To examine if the downstream target genes of the tumor suppressors p53 and Smad4 are affected by the sW182* mutant, we performed luciferase reporter assays. p21^Cip1^ is known to be regulated by both p53 and TGF-β/Smads [24]. The Smad7 promoter is a natural TGF-β/Smads-responsible promoter, and 3TP-lux is an artificial promoter for TGF-β/Smads [25,26]. Each of these promoter activities was inhibited by the sW182* mutant (Fig. 5A).

**Fig. 5.**
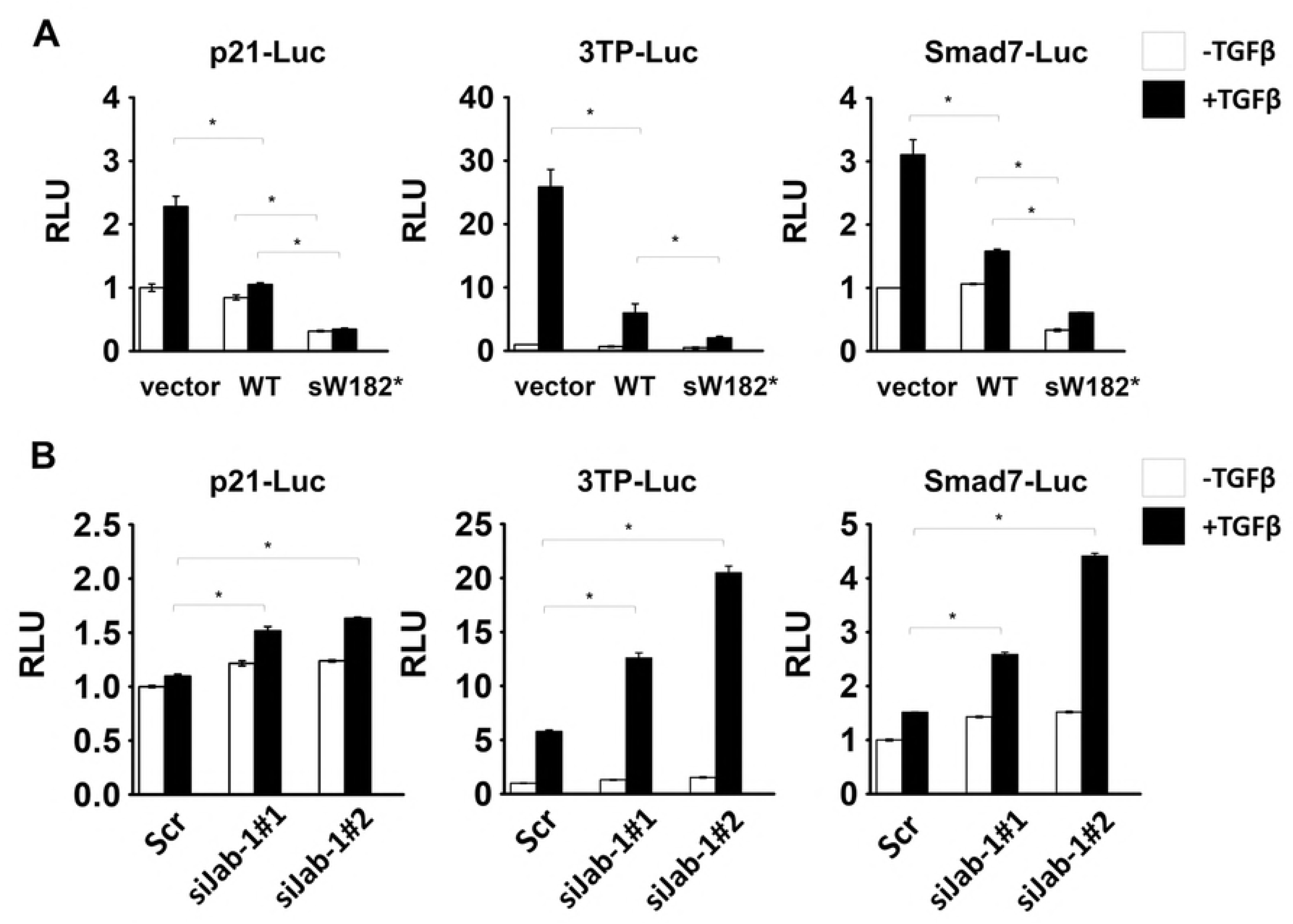
Effect of the LHBs sW182* mutant on downstream target genes of p53 and Smad4. (A) Huh-7 cells were cotransfected with the reporter construct along with vector, wild-type LHBs or sW182* mutant. Twenty-four h post transfection, the cells were treated with 0.5 nM TGF-β for another 24 h before the measurement of the luciferase activity. (B) The Huh-7 cells were transfected with siRNA. The siRNA-transfected cells were further transfected, treated with TGF-β and assayed for luciferase as in (A). Results shown are presented as means ± S.E.

This inhibition was relieved in the cells containing the Jab1 siRNA (Fig. 5B). These results indicated that the Jab1-dependent downregulation of the tumor suppressors by the LHBs sW182* mutant can influence their downstream target genes.

## Discussion

We and Lee et al. have recently identified nonsense mutations of the LHBs gene from the HBV-related HCC liver tumor tissues containing freely replicating HBV (HBV core antigen-positive, HBcAg(+)) [18,19]. These tumors with the freely replicating HBV were associated with significantly small sizes compared with those without detectable virus (HBcAg(-)), suggesting that they were in the early stages of tumorigenesis [19]. The presence of the nonsense mutants in early stage tumors also suggested that they might be able to play a direct role in hepatocarcinogenesis. This idea was tested in vitro and in vivo using mouse fibroblasts, NHI3T3. The three nonsense mutants, designated sL95*, sW182* and sL216*, have been shown to have oncogenic property and sW182* was the most potent [19]. In NIH3T3 cells, the sW182* elevated pro-oncogenic signals that include JNK, JAK1,2 and STAT3 [19]. The sW182* mutant can also epigenetically affects gene expression [20]. These effects may account in part for the sW182* mutant-induced tumorigenesis in the mouse xenograft model. These observations are, however, based on the transfected fibroblast cell line that is unrelated to the liver. For the functional studies it would be more relevant to use a cell line of liver origin. In the current study, we therefore examine the effects of the LHBs sW182* mutant in a liver cell line. Because of the labile nature of the mutant in the cultured liver cells, we only transiently expressed the sW182* in Huh-7 cells, which showed a low, but reasonable amount of the protein expression (Fig. 1).

The instability of the sW182* mutant in the cultured liver cells raised a concern about its functional role in liver tumorigenesis. The expression of the sW182* mutant in the mouse liver in vivo resolved the concern, as we confirmed a prolonged expression of the mutant protein in the liver cells in the living mice (Fig. 2). The in vivo expression of the wild-type and mutant proteins revealed a significant difference of the conditions surrounding the viral proteins between in vitro and in vivo. In the hydrodynamically injected mouse liver, the wild-type LHBs was hardly detected. This is not because of a failed expression, but rather secretion of the protein out from the liver. This idea was supported by detection of a high level of the wild-type LHBs in the blood samples. In contrast, the sW182* mutant was not detected in the blood but retained in the liver at a relatively high level. This nature of the truncation mutant would have a significant advantage for the induction of hepatocarcinogenesis if it has an oncogenic potential in the liver cells. The wild-type LHBs does not have this advantage.

Although the in vivo transfection by hydrodynamic injection showed a relatively strong and prolonged expression of the LHBs sW182* protein in the mouse liver, this method is considered as transient transfection with a limited efficiency (only part of the liver cells are transfected). To examine the functional effect of the LHBs sW182* protein on the mouse liver, a transgenic mouse model would be necessary.

From the observations above, we reasoned that mechanistic analysis of the sW182* mutant in the cultured liver cells is relevant, although it is unclear at present what causes the mutant protein unstable in the cultured liver cells. It has been reported that another mutant of LHBs protein (Pre-S2 mutant) can bind to Jab1 [16]. This study revealed that the LHBs protein contains a potential binding site for Jab1. This binding site appears to be hidden in the wild-type LHBs as it cannot bind to Jab1 (Fig. 3) [16]. When truncated, the binding site becomes available and associates with Jab1 (Fig. 3). This interaction is important for downregulation of downstream target tumor suppressor proteins.

We observed two major tumor suppressors p53 and Smad4, and their downstream target genes, were downregulated by the LHBs sW182* mutant, but not by the wild-type protein (Figs. 4,5). The downregulation of these tumor suppressors is, at least in part, mediated by proteasome as a proteasome inhibitor inhibits it (Fig. 4). These proteins are among the targets of Jab1 for degradation [30-33]. Consistently, depletion of Jab1 from the liver cells impaired the effect of the LHBs sW182* to downregulate the tumor suppressors and their target genes (Figs. 4,5). Our study strongly supports a Jab1-mediated pathway leading to tumor suppressor downregulation initiated by the truncated LHBs mutant. Jab1 can affect a number of other proteins positively or negatively [20]. Among them p27^Kip1^ is well known target of Jab1 [34,35]. It would be worthwhile to analyze more putative Jab1 targets if any of them is affected to further understand the signaling pathways that are influenced by the LHBs sW182* mutant. Such information would be very useful for development of novel therapeutic strategies for certain types of HCC.

In conclusion, the current study identified that the oncogenic LHBs truncation mutant sW182* downregulates multiple tumor suppressors through a Jab1-mediated pathway in the liver cells. The function of the truncation mutant would be enhanced by its strong liver-tropism. The downregulation of p53 and Smad4 can contribute the hepatocarcinogenesis in the HBV-infected liver containing the truncated viral protein.

## Acknowledgements

We are grateful to Mr. Shih-Hsuan Chan for his assistance in preparation of graphics.

